# IBD risk locus rs1077773 is a pharmacogenomic eQTL for aryl hydrocarbon receptor activity and modulates immune cell function

**DOI:** 10.1101/2025.09.26.678301

**Authors:** Ashley C. King, Kristen Seiler, Kerry Swanson, Matthew A. Ciorba, David M. Alvarado

## Abstract

**Introduction:** The inflammatory bowel diseases (IBD) Crohn’s disease (CD) and ulcerative colitis (UC) are disorders that cause chronic inflammation of the gastrointestinal tract. Both genetic and environmental factors contribute to the pathogenesis of IBD. There are currently >200 known genetic susceptibility loci for the development of IBD. The physiological impact of the majority of these loci remain a gap in our knowledge. One such locus is the single nucleotide polymorphism rs1077773, located ∼56kbp downstream from the aryl hydrocarbon receptor (*AHR*) gene. AHR is a ligand-activated transcription factor that is crucial to maintaining intestinal homeostasis. We hypothesized that rs1077773 enhances AHR activity to regulate mucosal immune response and maintain intestinal homeostasis.

**Methods:** All study procedures and reagents were approved by the Washington University Institutional Review Board (#202011003). Patient biopsies were collected at Barnes Jewish Hospital and genotyped using the IBD Genetics Consortium custom GSA SNP chip (Broad Institute) followed by imputation using TopMed Imputation Server at University of Michigan. Patient derived organoids (PDOs; N=3 G/G, N=4 G/A, N=5 A/A) were derived and maintained in 3D culture and supplemented with 50% L-WRN conditioned medium with passage every 3-4 days as previously described. PDOs were treated with AHR agonist 6-Formylindolo[3,2-b]carbazole (FICZ) or vehicle for 48h. Expression of *AHR* and its transcriptional targets Cytochrome P450 1A1 (*CYP1A1*) and *CYP1B1* was assessed by RT-qPCR. Blood was collected from pediatric patients undergoing intestinal resection at St. Louis Children’s Hospital and was genotyped with custom TaqMan SNP assay (N=3 G/G, N=5 G/A). Peripheral blood monocyte-derived macrophages (MDMΦs) were treated with lipopolysaccharide in the presence or absence of AHR ligands FICZ or indole-3-carboxaldehyde for 24h. Cytokine levels in culture supernatant were measured via using the ProcartaPlex human cytokine, chemokine, and growth factor 45-plex (ThermoFisher) on a Luminex FLEXMAP3D instrument.

**Results:** *AHR* expression was similar across genotypes and treatments. PDOs homozygous for rs1077773 demonstrate enhanced *CYP1A1* expression in response to AHR activation. In PBMΦs, cytokine secretion was stimulated by LPS treatment and was abrogated by FICZ treatment. PBMΦs with rs1077773 alternate allele demonstrated significant reduction in secretion of 17 cytokines and chemokines.

**Conclusions:** This work demonstrates that rs1077773 is an expression quantitative trait locus (eQTL) for AHR activity and modulates epithelial and immune cell function in vitro. Further mechanistic understanding of this locus and its correlates could improve our understanding of the molecular mechanisms of IBD susceptibility and may lead to novel personalized therapeutic approaches in IBD.

**Summary:** Our work demonstrates rs1077773 alternate allele is associated with enhanced aryl hydrocarbon transcriptional activity in human primary epithelial organoids and reduced lipopolysaccharide-induced inflammatory cytokine production in human peripheral blood monocyte-derived macrophages.

**Key Message:** We have identified rs1077773 as a pharmacogenomic regulator of inflammatory response in human primary intestinal organoids and peripheral blood monocyte-derived macrophages *via* AHR activity. Individual genetic variation affecting this pathway may account for differences in response to environmental stimuli and the development and progression of IBD.

## Introduction

The intestinal epithelium forms a physical, chemical, and immunological barrier between an individual and their external environment, including the intestinal microbiome. Chronic disruptions in this barrier, dysregulated inflammatory responses, and dysbiosis of the intestinal microbiome are hallmarks of inflammatory bowel diseases (IBD) Crohn’s disease (CD) and ulcerative colitis (UC). IBD arises through a combination of genetic and environmental factors.^1^ Genome-wide association studies (GWAS) have identified hundreds of genetic susceptibility loci for IBD. While many loci have been mapped to functional changes in proteins involved in epithelial barrier function and immune regulation, the cellular impact of most genetic loci remains unknown. A better understanding of the genetic factors underlying epithelial barrier dysfunction in IBD could uncover novel diagnostic and therapeutic approaches for these disorders.

The aryl hydrocarbon receptor (AHR) is an environmental sensor that is activated by a variety of endogenous and exogenous stimuli and helps maintain barrier integrity and mucosal immune tolerance.^2^ The single nucleotide polymorphism (SNP) rs1077773:G>A was identified by GWAS as associated with reduced risk of IBD (odds ratio 0.93).^3^ This SNP is approximately 56 kilobase pairs 3’ from the *AHR* gene and is located in a genomic region predicted to modulate AHR promoter activity.^4^ SNP rs1077773 is among 12 loci predictive of durable response to anti-TNF therapy in UC patients, suggesting this region can modulate response to IBD therapies.^5^ However, the impact of rs1077773 on AHR activity and cellular function is not known. The present study uses primary intestinal patient-derived organoids (PDOs) and peripheral blood monocyte-derived macrophages (MDMΦs) to assess the physiological impact of rs1077773 at baseline and in response to AHR agonists.

## Methods

All study procedures and reagents were approved by the Washington University Institutional Review Board (#202011003). Patient demographics for PDOs and MDMΦs are shown in **Table 1**. PDOs were genotyped using an Illumina custom global screening array developed by the IBD genetics consortium (IBDGC).^6^ Data were provided as pre-processed binary PLINK files and SNP rs1077773 was imputed through TopMed Imputation Server at University of Michigan (r^2^>0.3).^7^ PB-mΦ donors were genotyped using TaqMan® custom assay (ANPR43G, ThermoFisher Scientific).

**Table 1:** Donor demographics for patient-derived intestinal organoids (PDO) and peripheral monocyte-derived macrophages (MDMΦs). G/G, rs1077773 homozygous reference allele; G/A, heterozygous; A/A, homozygous alternate allele; CD, Crohn’s disease; UC, ulcerative colitis; HD, Hirschsprung’s disease; IND, indeterminate colitis; SD, standard deviation; Categorical data were compared using Chi square test, and continuous data were compared using student’s t test/ANOVA.

| PDO cohort |  | G/G | G/A | A/A | P value |
| --- | --- | --- | --- | --- | --- |
| Diagnosis | None | 1 (33.3) | 0 (0) | 3 (60) | 0.2598 |
|  | CD | 2 (66.7) | 2 (50) | 1 (20) |  |
|  | UC | 0 (0) | 2 (50) | 1 (20) |  |
| Age | Mean (SD) | 44.7<br>(5.31) | 47.2<br>(16.8) | 48.6<br>(13.0) | 0.72 |
| Sex | Female (%) | 2 (66.7) | 2 (50) | 3 (60) | 0.9134 |
|  | Male (%) | 1 (33.3) | 2 (50) | 2 (40) |  |
| Race | White (%) | 2 (66.7) | 5 (100) | 4 (80) | 0.7732 |
|  | Black (%) | 1 (33.3) | 0 (0) | 1 (20) |  |
|  | Jewish (%) | 0 (0) | 1 (20) | 0 (0) |  |

| PB-mΦ cohort |  | G/G | G/A | P value |
| --- | --- | --- | --- | --- |
| Diagnosis | None | 1 (33.3) | 1 (20) | 0.5541 |
|  | CD | 2 (66.7) | 1 (20) |  |
|  | UC | 0 (0) | 1 (20) |  |
|  | HD | 0 (0) | 1 (20) |  |
|  | IND | 0 (0) | 1 (20) |  |
| Age | Mean (SD) | 10.4 (7.2) | 9.3 (6.1) | 0.864 |
| Sex | Female (%) | 1 (33.3) | 2 (40) | 0.850 |
|  | Male (%) | 2 (66.7) | 3 (60) |  |

PDOs were derived and maintained as previously described.^8^ Briefly, cells were embedded in Matrigel and supplemented with 50% L-WRN conditioned medium with passage every 3-4 days. Cells were incubated with 1nM 6-formylindolo[3,2-b]carbazole (FICZ; Sigma) or vector (DMSO) for 48h before RNA isolation with Qiagen RNEasy Mini kit. RNA expression was assessed using real-time qPCR with the following primers: *ACTB* (Fwd ATCATTGCTCCTCCTGAGCG; Rev GCTGATCCACATCTGCTGGAA), *AHR* (Fwd ACATCACCTACGCCAGTCG; Rev CGCTTGGAAGGATTTGACTTGA), *CYP1A1* (Fwd ACATGCTGACCCTGGGAAAG; Rev GGTGTGGAGCCAATTCGGAT), *CYP1B1* (Fwd AAGTTCTTGAGGCACTGCGAA; Rev GGCCGGTACGTTCTCCAAAT) and *GAPDH* (Fwd GACCTGCCGTCTAGAAAAACC; Rev GCTGTAGCCAAATTCGTTGTC).

PBMCs were isolated from whole blood via density separation with Ficoll-Paque (ThermoFisher Scientific) and monocytes were isolated by adherence to cell culture plates for 1h in complete medium (RPMI 1640 medium supplemented with 10% FBS and 1% penicillin/streptomycin).^9^ Monocytes were differentiated into peripheral blood-derived macrophages (MDMΦs) over 6 days in complete medium supplemented with 50ng/ml macrophage colony stimulating factor (M-CSF). MDMΦs were treated 24h in complete medium supplemented with 50ng/ml M-CSF alone (CTRL), with 10ng/ml lipopolysaccharide (LPS), with LPS and 10nM FICZ (LPS + FICZ), or LPS and 100uM indole-3-carboxaldehyde (LPS + I3CA). Cytokines were quantified in culture supernatant using the ProcartaPlex human cytokine, chemokine, and growth factor 45-plex (ThermoFisher) on a Luminex FLEXMAP3D instrument (Luminex Corporation, Austin, TX). The standard curve was generated using a 5-parameter curve fit and Belysa software v1.2.1 (MilliporeSigma) and the sample mean fluorescent intensities (MFI) for fifty beads per region were quantified.

Data analysis was performed using RStudio 2023.06.0+421 “Mountain Hydrangea” Release (583b465e, 2023-06-05) for windows. PCR results were normally distributed (Wilke’s Normality test) and were analyzed using one-way ANOVA. Cytokine quantification results were subjected to linear regression analysis with experimental batch effects and patient demographics (age, sex, race, and diagnosis) included as covariates. P<0.05 was considered significant.

## Results

To assess the physiological impact of rs1077773 in intestinal epithelial cells, we cultured PDOs with AHR ligand FICZ or without (CTRL) and total RNA was collected after 48h. Expression of *AHR* and its target genes *CYP1A1* and *CYP1B1* was assessed by quantitative realtime PCR. *AHR* expression was not affected by rs1077773 genotype at baseline or following FICZ treatment (**Fig. 1A**). However, the rs1077773 alternate allele was associated with significantly higher *CYP1A1* (P=0.0257 by one-way ANOVA; **Fig. 1A**) and a trend towards increased *CYP1B1* expression (**Fig. 1B**). To further explore the relationship of rs1077773 genotype to AHR activity, a series of generalized linear models were applied to the expression data with and without patient age, sex, race, and diagnosis as covariates. *AHR* expression was not affected by any factor, although patient sex had the most impact (AHR ∼ Geno_b + Age + Sex + Race + Diagnosis, P=0.403 overall, P=0.0525 sex). Patient demographics did not impact *CYP1A1* expression independently (CYP1A1 ∼ age + sex + race + diagnosis, P=0.6358) or collectively (CYP1A1 ∼ Age:Sex:Race:Diagnosis, P=0.6079). In contrast, *CYP1A1* expression was significantly altered by rs1077773 genotype, with the alternate allele associated with higher *CYP1A1* expression (*CYP1A1* ∼ genotype, P=0.02248). One datapoint for *CYP1B1* expression was greater than 2 standard deviations from the mean, which defines this value as a statistical outlier (P=0.01). None of the linear models demonstrated significance when this outlier was included in analysis. Exclusion of this sample revealed rs1077773 genotype as the only factor impacting *CYP1B1* expression (CYP1B1 ∼ genotype + age:sex:race:diagnosis, P= 0.003753). These data demonstrate that rs1077773 modulates the activity of AHR in a genotype- and ligand-dependent, or pharmacogenomic, manner in intestinal epithelial cells.

**Figure 1:**
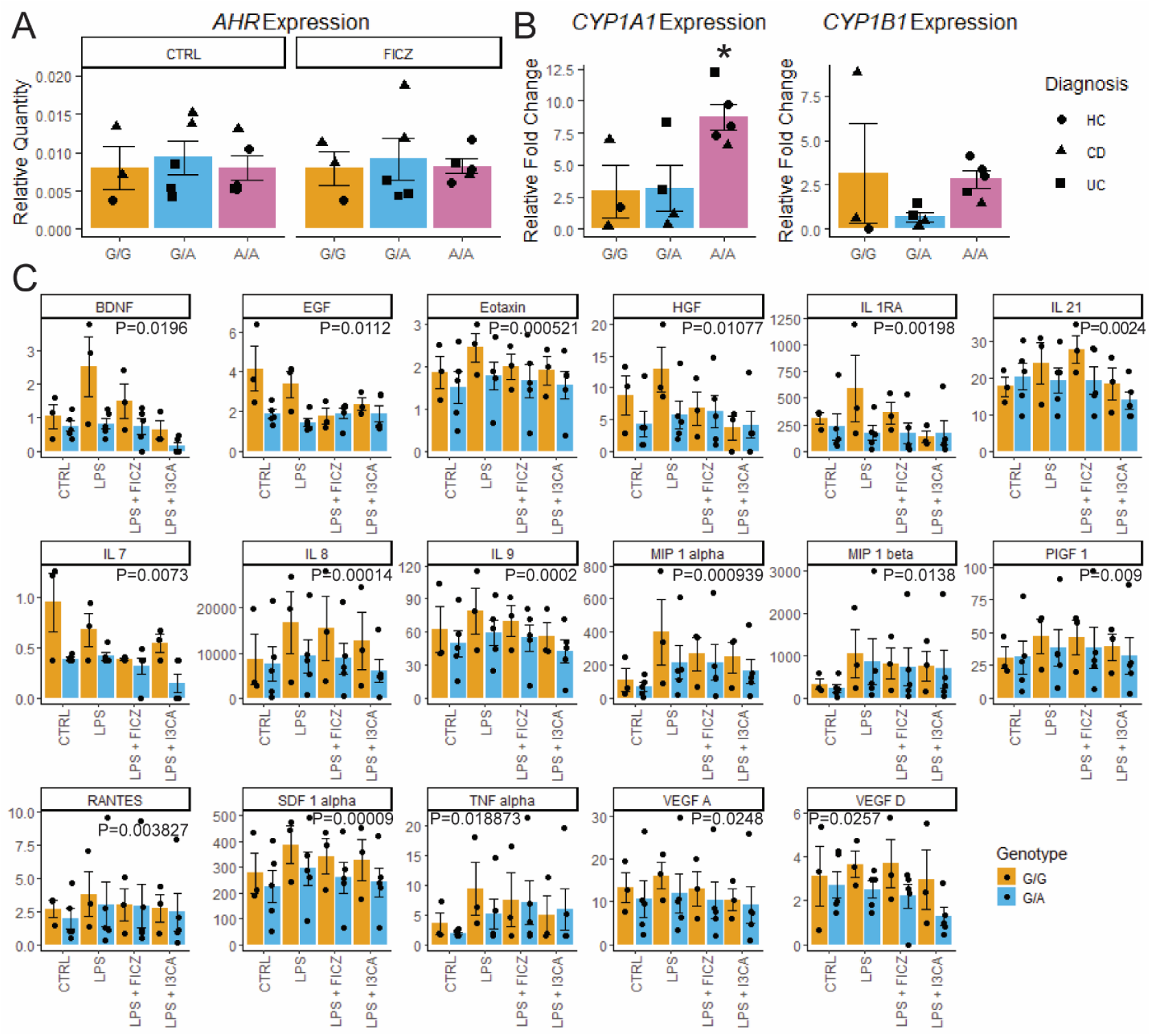
SNP rs1077773 modulates AHR activity and MDMΦs function. A) Quantification of *AHR* RNA expression relative to *ACTB/GAPDH*. B) Relative fold change of AHR targets *CYP1A1* and *CYP1B1* in response to AHR ligand FICZ. *P=0.02571 for *CYP1A1* ∼ SNP allele by linear regression model. C) Concentration (pg/ml) of 17 cytokines which demonstrate significant reduction in heterozygous MDMΦs (G/A) compared to homozygous reference allele (G/G).

Next, MDMΦs were derived from 8 pediatric patients undergoing surgical resection at Saint Louis Children’s Hospital (**Table 1**). These cells were treated with vehicle (CTRL), LPS, LPS + FICZ, or LPS + AHR ligand I3CA for 24h. Cytokine levels were assessed in the culture supernatant using a Luminex FLEXMAP3D instrument. Three of the 45 cytokines (IL-12p70, Ifn-α, Ifn-γ) were only detectable in a minority of samples and were excluded from analysis. The remaining data were analyzed by linear regression models examining the association of cytokine levels to treatment and rs1077773 genotype. Due to limited sample size, patient diagnosis was excluded from analysis. After adjusting for age, sex, and batch effects, 17 cytokines demonstrated significantly reduced cytokine levels associated with the alternative allele (**Fig. 1C**). In most cases, LPS treatment resulted in increased cytokine levels, while treatment with AHR ligands reduced these cytokine levels. Six cytokines were significantly modulated by at least one treatment but not by rs1077773 genotype (IL-13, IL-17A, IL-2. IL-23, IL-5, and PDGF BB), while the remaining 19 cytokines did not vary significantly across any treatment. Taken together, these data demonstrate rs1077773 impacts cytokine secretion in MDMΦs, with the alternate allele associated with reduced cytokine levels.

## Discussion

Genetic predisposition and environmental exposures contribute to the development of IBD, though a clear molecular mechanism is not known for many genetic susceptibility loci.^1^ In the present study we show that rs1077773 modulates AHR transcriptional activity and cell function in response to inflammatory stimulus. To our knowledge, this is the first study to demonstrate a cellular phenotype associated with rs1077773. These findings suggest altered AHR pathway activity could contribute to physiological differences that modulate IBD susceptibility. However, the molecular mechanism remains unclear. Our study does not show differences in the expression of *AHR* associated with rs1077773. This is surprising, as a previous study demonstrated the genomic region inclusive of rs1077773 is capable of driving transcription of a luciferase reporter *in vitro*.^4^ However, these data do not preclude indirect mechanisms of transcriptional regulation, such as modulation of chromatin accessibility or other long-range genomic interactions. Further investigation into the mechanism(s) by which rs1077773 can modulate AHR pathway activity is needed.

This study illustrates how the interaction of individual genetic variation and environmental factors that impact AHR activity could contribute to the development of IBD. These pharmacogenomic interactions may also impact response to IBD therapies, as rs1077773 was one of 12 genetic markers that predict durable response to anti-TNF therapy in UC patients.^5^

Furthermore, AHR is the target for the herbal extract *Indigo naturalis*, which is currently being explored in combination with curcumin as a therapeutic agent for UC with promising results.^10^ Thus, individual genetic variation impacting the AHR pathway may be important to consider in therapy selection, paving the way towards personalized medicine in IBD. Some limitations of this work include the small sample size and the lack of individuals homozygous for the alternate allele in the pediatric surgery cohort. A more comprehensive assessment of individual genetic variation affecting AHR signaling is needed to fully delineate the role of AHR pharmacogenomics in the maintenance of intestinal homeostasis, the development and progression of IBD, and response to therapies.

## Abbreviations

AHR: Aryl hydrocarbon receptor
MDMΦs: Monocyte-derived macrophages
PBMCs: Peripheral blood mononuclear cells
IBD: inflammatory bowel diseases
CD: Crohn’s disease
UC: Ulcerative colitis
PDOs: Patient-derived organoids

## Acknowledgments

We thank the Washington University Digestive Disease Research Core Center for access to patient samples and cell culture support (P30 DK052574). This project was supported by the Washington University Institute of Clinical and Translational Sciences (ICTS) which is, in part, supported by the NIH/National Center for Advancing Translational Sciences (NCATS), CTSA grant #UL1TR002345.

We thank Dr. Judy Cho at Mount Sinai Ichan School of Medicine for assistance with SNP genotyping. Global screening array was performed at the Feinstein Institute for Medical Research in conjunction with the National Institute of Diabetes and Digestive and Kidney Diseases (NIDDK) IBD Genetics Consortium (IBDGC; UM1 HG008901). This manuscript was not prepared in collaboration with the IBDGC and does not necessarily reflect the opinions or views of the IBDGC investigators or the NIDDK.

## Notes

**Support** NIH R01-AI167285 (MAC); Crohn’s and Colitis Foundation #648423 (DMA) and the Washington University Institute of Clinical and Translational Sciences (ICTS) which is, in part, supported by the NIH/National Center for Advancing Translational Sciences (NCATS), CTSA grant #UL1TR002345; This work was supported, in part, by the Bursky Center for Human Immunology and Immunotherapy Programs at Washington University, Immunomonitoring Laboratory.

**Authors’ financial disclosures** The authors have no financial disclosures or conflicts of interest.

### Competing Interest Statement

The authors have declared no competing interest.

